# A VPSl5-like kinase regulates apicoplast biogenesis and autophagy by promoting PI3P generation in *Toxoplasma gondii*

**DOI:** 10.1101/2022.05.30.493977

**Authors:** Rahul Singh Rawat, Priyanka Bansal, Pushkar Sharma

## Abstract

Phosphoinositides are important second messengers that regulate key cellular processes in eukaryotes. While it is know that a single phosphoinositol-3 kinase (PI3K) catalyses the formation of 3’-phosphorylated phosphoinositides (PIPs) in apicomplexan parasites *Plasmodium* and *Toxoplasma,* how its activity and PI3P formation is regulated has remained unknown. Present studies involving a unique Vps15 like protein (TgVPS15) in *Toxoplasma gondii* provide insights into the regulation of phosphatidyl-3-phosphate (PI3P) generation and unravel a novel pathway that regulates parasite development. Detailed investigations suggested that TgVPS15 regulates PI3P formation in *Toxoplasma gondii,* which is important for the inheritance of the apicoplast-a plastid like organelle present in most apicomplexans and parasite replication. Interestingly, TgVPS15 also regulates autophagy in *T. gondii* under nutrient-limiting conditions as it promotes autophagosome formation. For both these processes, TgVPS15 uses PI3P-binding protein TgATG18 and regulates trafficking and conjugation of TgATG8 to the apicoplast and autophagosomes, which is important for biogenesis of these organelles. TgVPS15 has a protein kinase domain but lacks several key residues conserved in conventional protein kinases. Interestingly, two critical residues in its active site are important for PI3P formation and parasitic functions of this kinase. Collectively, these studies unravel a signalling cascade involving TgVPS15, a novel effector of PI3-kinase in *T. gondii* and possibly other Apicomplexa, that regulate critical processes like apicoplast biogenesis and autophagy.

## Introduction

*Toxoplasma gondii* is an obligate intracellular protozoan and belongs to the phylum Apicomplexa. It shares several features with *Plasmodium* spp., which cause malaria and is considered as an excellent model to study obligate parasitism. Most apicomplexan parasites possess an essential plastid-like organelle called apicoplast, which is surrounded by four membranes, acquired as a result of two successive endosymbiosis events (McFadden et al, 1996). The apicoplast segregates between daughter cells during cell division and it harbours key biochemical pathways, which makes it essential for the parasite survival (Roos et al, 1999; Striepen et al, 2007).

Phosphoinositides (PIPs) are important second messengers that regulate key processes in eukaryotes and are generated by the action of phosphoinositide kinases (PIKs) on precursor PIPs (Balla, 2013). Of late, there has been an upsurge in interest in PIP signaling in apicomplxean parasites such as *Toxoplasma* and *Plasmodium* spp. as they have been implicated in several critical processes like organelle biogenesis, protein export and trafficking, host cell invasion (Wengelnik et al, 2018). PI3-kinases (PI3Ks) belong to three major classes and class III PI3Ks are mainly involved in the generation of PI3P (Vanhaesebroeck et al, 2001). *Toxoplasma* as well as *Plasmodium* code for a single PI3K, which closely resembles class III PI3Ks and generates PI3P in these parasites (Daher et al, 2015; Vaid et al, 2010). TgPI3K/PI3P regulates biogenesis of apicoplast in *Toxoplasma gondii* (Daher et al, 2015). PI3P typically executes its cellular functions via specific PI3P-binding domain containing proteins like FYVE or PX domains (Lemmon, 2003). Several PIP binding proteins have been identified in apicomplexa but function of most of these has remained unknown (Wengelnik et al, 2018). Recently, we demonstrated that autophagy related protein ATG18, which interacts with PI3P, is critical for apicoplast biogensis (Bansal et al, 2017). Autophagy related protein 8 (ATG8), which is localized to the apicoplast, in steady state conditions is indispensable for apicoplast biogenesis (Leveque et al, 2015). These proteins are typically associated with autophagy in yeast and mammals. Autophagy is a catabolic process used by most eukaryotic cells to discard damaged and unwanted material as well as recycle key components for their survival under nutrient-limiting conditions (Feng et al, 2014). The formation of autophagosome is critical for the sequestration and delivery of cellular material to the lysosome for degradation. The process of autophagy is a tightly regulated process involving the formation of specific protein complexes and conjugation of ATG8 to the autophagosome membrane is critical for this process. PI3P plays a critical role in autophagy (Burman & Ktistakis, 2010). Most eukaryotes possess core autophagy machinery and this machinery seems to be simpler in *Apicomplexa.* However, upstream components are either divergent or are absent (Feng et al, 2014; Sakamoto et al, 2021). ATG8 and proteins/enzymes involved in its processing are largely conserved in *Toxoplasma* and other apicomplexans and also contribute to autophagy (Kong-Hap et al, 2013).

In our efforts to understand the mechanisms involved in the generation of PIPs, which are not understood in apicomplexan parasites, we investigated the role of Vps15 like protein from *Toxoplasma.* Vps15 has been reported to increase the activity of Vps34 like class III PI3-kinases in yeast and other organisms (Stack & Emr, 1994). While it is classified as a “pseudokinase”, the yeast homologue has been reported to exhibit autophosphorylation (Stack & Emr, 1994) but no substrates of this kinase have been reported. However, the human homologue p150 does not exhibit autophosphorylation (Panaretou et al, 1997). While it is unclear if it regulates PI3Ks by phosphorylation, Vps15 is part of two major multiprotein complexes comprising of Vps34 (PI3K) and several other proteins like ATG14L, Beclin1 in Complex I and VPS38/UVRAG replaces ATG14 in Complex II. Complex I is important for autophagosome formation as it promotes PI3P generation at the phagophore. Complex II regulates various other cellular processes like endosome sorting, lysosome recycling, autophagosome maturation and cytokinesis (Ohashi et al, 2019). Recent structural analysis of Complex II shed light on the organization of the complex as well as provides important details related to the regulation of individual protein components like VPS15 (Rostislavleva et al, 2015).

Apicomplexan parasites *Plasmodium falciparum* and *Toxoplasma gondii* code for a VPS15 orthologue with interesting differences, such as domain organization and size. In the present study, we demonstrate that *Toxoplasma gondii* homologue TgVPS15 facilitates PI3P formation and regulates apicoplast inheritance as well as autophagy under starvation conditions.

## Results

### A Vps15 orthologue in Plasmodium and Toxoplasma

We were interested in understanding the regulation of PI3P in apicomplexan parasites *Toxoplasma* and *Plasmodium*. In our quest to identify putative Vps15 homologues, *in silico* sequence based search revealed VPS15 orthologues in *Toxoplasma gondii* (ToxoDB ID: TGGT1_310190) and a *P. falciparum* (PlasmoDB ID: PF3D7_0823000). These observations were consistent with previous studies, which also indicated the presence of these genes as putative VPS15 orthologues in these parasites (Navale et al, 2014) (Smith et al, 2021). However, VPS15 has not been analyzed and characterized and its function in parasite biology has remained unknown. The yeast ScVps15 has a kinase domain at the N-terminus followed by Armadillo like helical repeats (HEAT) in the middle that are connected to a six WD40 repeats at the C-terminus, which form a β-propeller structure (Backer, 2016; Rostislavleva et al, 2015). The domain analysis of TgVPS15 suggested that it possesses the kinase and the Helical or HEAT domain (Fig. 1A). However, various algorithms could at best predict two highly degenerate WD40 repeats with very low scores in the case of TgVPS15. Therefore, it is possible that TgVPS15 does not contain WD40 repeats. Alternatively, it is possible that the corresponding C-terminal region may lack sequence homology but may fold into β-propeller structure observed in the yeast VPS15. Detailed structural analysis will only be able to test these possibilities. In the case of PfVPS15, the WD40 repeats could not be predicted with certainty. Moreover, PfVPS15 is a much shorter protein and ends with the Helical or HEAT domain (Fig. 1A). These observations suggested that VPS15 in these parasites has a highly divergent domain arrangement when compared to the yeast and mammalian counterparts.

**Figure 1.**
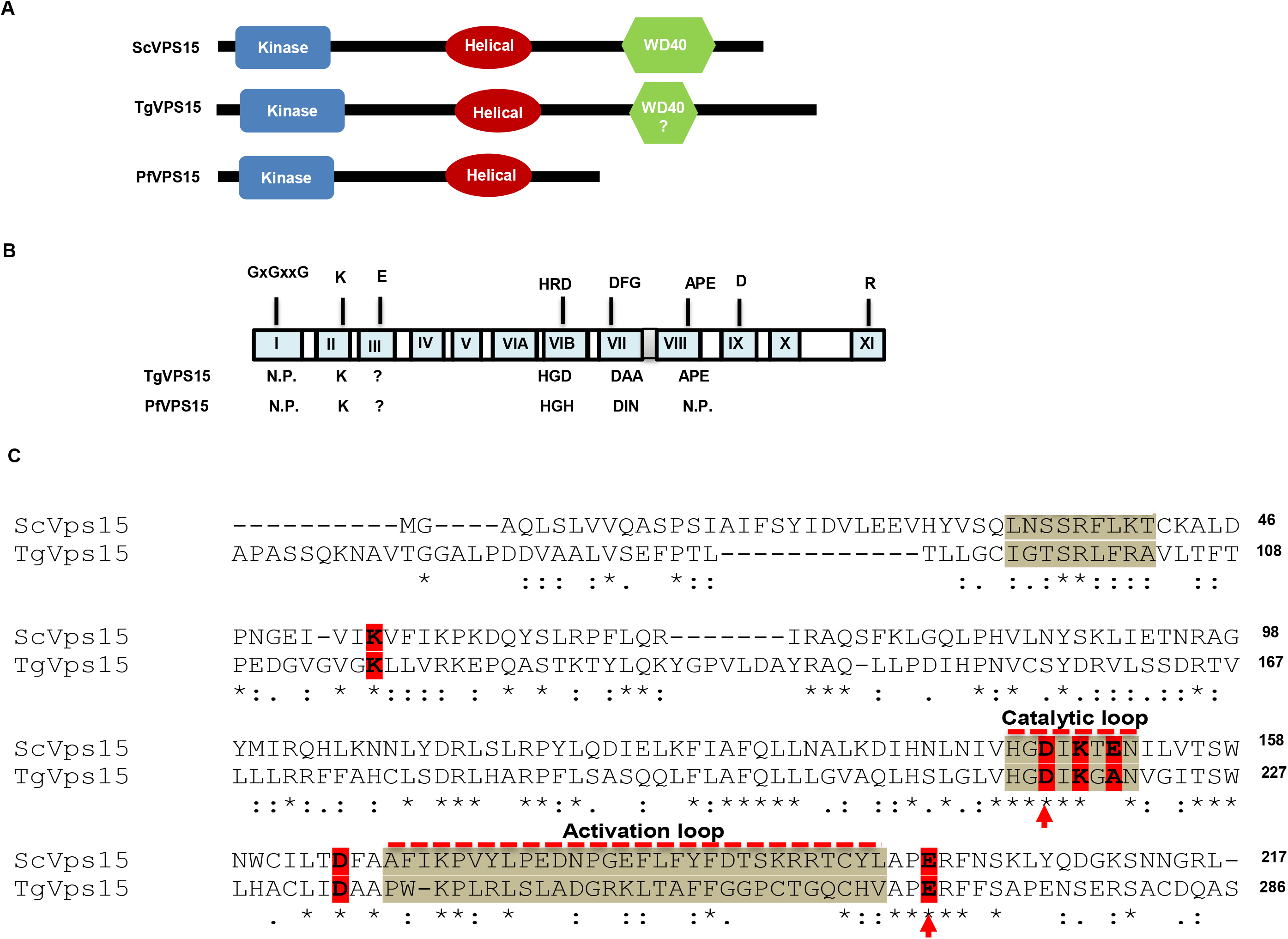
A VPS15 orthologue in *P. falciparum* and *Toxoplasma gondii*. **A**. Domain organization of Pf/TgVPS15 revealed that TgVPS15 contains a kinase like domain (KD) at the N-terminus followed by helical HEAT repeats, which are connected by long linkers. WD40 could not be predicted with certainty as only weak sequence similarity was found to WD40 repeats in the case of TgVPS15 whereas PfVPS15 does not contain the C-terminal WD40 domain. B. Schematic illustrating the subdomains of canonical protein kinases and key putative residues involved in catalysis are indicated on the top and corresponding residues in Tg/PfVPS15 are indicated below. N.P. - not present. C. CLUSTALW alignment of the kinase domains of ScVPS15 with TgVPS15. Some of the key motifs and subdomains are indicated and putative residues possibly involved in kinase function are highlighted. D216 and E268, which were mutated in subsequent studies, are indicated by an arrow.

VPS15 from yeast and mammals has been classified as a pseudokinase as it lacks key catalytic residues present in protein kinases (Fig. 1B) (Herman et al, 1991; Stack et al, 1993). We analyzed the sequence of the kinase domain of Tg/PfVPS15 by comparing it with that of ScVPS15. Like several pseudokinases and ScVps15 (Rostislavleva et al, 2015), TgVPS15 lacks key motifs: a. GxGxxG motif in the ATP binding loop is absent in both ScVPS15 and TgVPS15; b. Arginine in the HRD motif of the catalytic loop, which is important for the stabilization of activation loop of kinases that need to undergo autophosphorylaion of this loop for activation (Johnson et al, 1996; Nolen et al, 2004) is also missing from both ScVPS15 and TgVPS15. However, the catalytic aspartate D216 in TgVPS15 is conserved in these kinases; c. The aspartate in the DFG motif of the catalytic loop of protein kinases is critical for chelating divalent cations like Mn or Mg and is conserved in TgVPS15 (D234) and ScVPS15, whereas the following phenylalanine is absent in TgVPS15 (Fig. 1C); c. APE motif, which is part of P+1 loop (Nolen et al, 2004), is well conserved in these kinases. The activation loop of protein kinases resides between the DFG (aa 234-236 in TgVPS15) and APE (aa 266-268 in TgVPS15) and is critical for their activation and often undergoes autophosphorylation in several conventional kinases especially the ones with RD motif (Nolen et al, 2004). While the activation loop is present, it seems to show only marginal conservation between ScVPS15 and TgVPS15. Interestingly, sequence comparison suggests that PfVPS15 may lack the activation loop (not shown here).

### TgVPS15 regulates PI3P formation in the parasite and its depletion impairs parasite replication

In order to investigate the function of TgVPS15, an inducible knockdown was generated in *T. gondii,* which was achieved by using a tetracycline regulated transactivator system (Salamun et al, 2014). A parasite line was generated in which the native promoter of TgVPS15 was exchanged with 7-tet-OpSag1 promoter (Bansal et al, 2021) and a Myc-tag was introduced at the N-terminus of the TgVPS15 5’-end and these genetic modifications were confirmed by genotyping (Supp. Fig. S1A, S1B).

The addition of ATc to TgVPS15-iKD parasites revealed depletion of TgVPS15 protein within 24h treatment as assessed by Western blotting (Fig. 2A) and also by immunofluoroscence assay (IFA) (Fig. 2B). IFA also revealed that TgVPS15 is present in punctate structures spread in the parasite cytoplasm (Fig. 2B).

**Figure 2.**
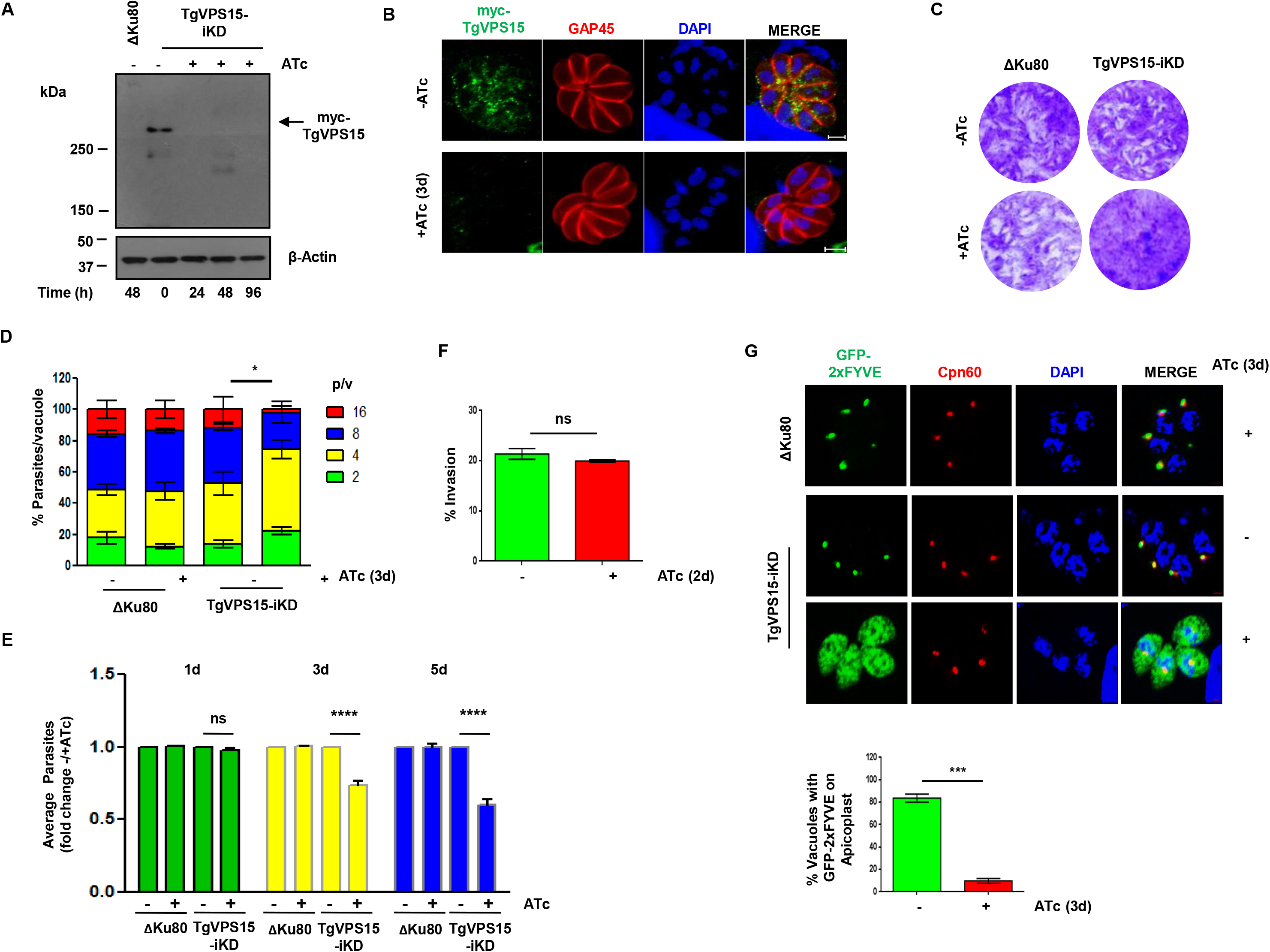
TgVPS15 regulates PI3P formation and is critical for parasite replication. A. Depletion of TgVPS15 in *Toxoplasma gondii.* TgVPS15-iKD parasites were treated with ATc for indicated duration followed by Western blotting of parasite lysate, which revealed that Myc-TgVPS15 migrates at an expected size and the addition of ATc resulted in its depletion. ΔKu80 was used as a negative control. B. IFA was performed on TgVPS15-iKD parasites that were left untreated or treated with ATc for 72h or 3d. using anti-myc and anti-GAP45 antibodies.. C. Plaque assays were carried out by infecting HFF monolayer with ΔKu80 or TgVPS15-iKD parasites in the presence or absence of ATc for 7 days (for quantitation please see Supp. Fig. S3B). D. TgVPS15-iKD or ΔKu80 parasites were preincubated in culture medium for 48h or 2d with (+) or without (-) ATc and were subsequently allowed to invade fresh HFFs in the presence or absence of ATc and the number of parasites per vacuole was determined after 24h or 1d. Data represent Mean ± SE, n=3 and at least 200 vacuoles were counted for each condition (n=3, *** P<0.001 4 p/v, ** P<0.01 8 p/v, * P<0.05 16 p/v ANOVA,, ns-not significant). E. TgVPS15-iKD or ΔKu80 parasites were treated with ATc for 1d, 3d or 5d and replication assay was performed as described in panel E and average intracellular parasites/vacuole present in each condition was determined. Fold change in average parasites upon ATc addition was determined (Mean ± SE, n=3, ANOVA, **** P<0.0001; ns -not signficant). F. Invasion assays were performed on TgVPS15-iKD parasites that were left untreated or treated with ATc. No significant change in invasion was observed (P> 0.05, t test, n=2, ns-not significant). G. ΔKu80/DD-GFP2xFYVE or TgVPS15-iKD/DD-GFP2xFYVE parasites that expressed GFP-2xFYVE domain infected HFF were left untreated (-) or treated (+) with ATc for 72h or 3d. Prior to fixation, Shield-1 was added for 30 minutes to stabilize the expression of GFP-2xFYVE, which was localized mainly to the apicoplast as revealed by IFA for Cpn60, an apicoplast protein. GFP-2xFYVE was found mainly in the cytoplasm of ATc-treated TgVPS15-iKD parasites. *Right Panel,* % vacuoles in which GFP-2xFYVE was found at the apicoplast was determined (Mean±SE, n=3, *** p<0.001, t-test).

TgVPS15-iKD parasites were used to investigate the function of TgVPS15 in parasite development. To monitor the parasite growth during the parasitic lytic cycle in host monolayer, plaque assays were performed that are indicative of crucial parasite processes like intracellular development/replication, host cell invasion and egress. ΔKu80 as well as TgVPS15-iKD parasites exhibited the formation of plaques (Fig. 2C, Supp. Fig. S3B). ATc treatment of TgVPS15-iKD parasites caused a marked reduction in plaques (Fig. 2C, Supp. Fig. S3B). Further studies were carried out to identify which developmental processes were regulated by TgVPS15. The effect of TgVPS15 depletion on parasite division was evaluated by performing intracellular parasite replication assays. There was no significant change in the average number of parasites after depletion of TgVPS15 in the first lytic cycle (24h/1d ATc addition) (Fig. 2E, Supp. Fig. S2). However, there was a marked difference in parasite replication in subsequent rounds of lytic cycle as 3 (Fig. 2D, 2E) or 5 days of ATc treatment (Fig. 2E)) lead to significantly fewer number of parasites per vacuole. To complement the TgVPS15 function in the TgVPS15-iKD line, a copy of *TgVPS15* was integrated into the TgVPS15-iKD genome and ectopically expressed using tubulin 1 promoter with an N-terminal Ty-tag (TgVPS15-iC^WT^) (Supp. Fig. S3A). Upon ATc treatment, the number of plaques formed in TgVPS15 complemented parasites (TgVPS15-iC^WT^) was not altered significantly (Supp. Fig. S3B). Furthermore, defects in parasite replication were reversed significantly in the case of TgVPS15-iC^WT^ parasites (Fig. 7B). These data suggested that TgVPS15 signaling and downstream events regulate replication of *T. gondii.* We also checked for the role of TgVPS15 in host cell invasion but there was almost no defect in invasion upon TgVPS15 depletion (Fig. 2F).

It was important to evaluate if TgVPS15 is involved in PI3P generation in the parasite. To this end, DD-GFP-2xFYVE reporter was ectopically expressed by integration in the TgVPS15-iKD parasites. The FYVE-reporter protein binds to PI3P via FYVE domain present in tandem repeats and has been previously reported to detect PI3P levels in *Toxoplasma* (Tawk et al, 2011). Since this reporter is fused to N-terminal FKBP-destabilization domain (DD) its expression can be regulated by the addition of Shld-1, which prevents its degradation by proteasomes. IFA studies suggested that FYVE domain was localized mainly to the apicoplast as indicated by co-localization of apicoplast resident protein Cpn60 (Fig. 2C), which was consistent with previous reports and suggested that apicoplast membrane is enriched in PI3P (Tawk et al, 2011). Upon TgVPS15 depletion, 2xFYVE was largely absent from the apicoplast and diffused in the cytoplasm, which indicated a significant reduction in PI3P at the apicoplast (Fig. 2D). These data strongly indicated that TgVPS15 is involved in PI3P generation at the apicoplast. Therefore, it is reasonable to suggest that TgVPS15 may regulate PI3-Kinase TgPI3K, which is responsible for catalyzing PI3P generation (Daher et al, 2015).

### TgVPS15 is involved in apicoplast inheritance

Our results suggested that defects in parasite replication were observed only in second and subsequent cycle after TgVPS15 depletion (Fig. 2 E). Such delayed death phenotype in apicomplexans like *Toxoplasma* and *Plasmodium* has been associated with defective parasite division after the first lytic cycle and is attributed to the loss of apicoplast (Fichera & Roos, 1997). Therefore, we analyzed the status of apicoplast formation, which is inherited from the mother to budding daughter cells during endodyogeny, by performing IFA for apicoplast resident protein Cpn60. Cpn60 is a nuclear encoded protein and contains a N-terminal apicoplast targeting signal, which is cleaved after its trafficking to the apicoplast. A significant loss of apicoplast was progressively observed after ATc treatment, which was indicated by either diffused or missing Cpn60 (Fig. 3A, B). Furthermore, the mature form of Cpn60-obtained after processing of its apicoplast targeting signal at the apicoplast was reduced and predominantly the unprocessed Cpn60 was observed in TgVPS15 depleted parasites (Fig. 3C). Consistent with these observations, apicoplast genome replication was also significantly impaired upon TgVPS15 depletion (Fig. 3D). Collectively, these data established a role of TgVPS15 in apicoplast biogenesis or inheritance.

**Figure 3.**
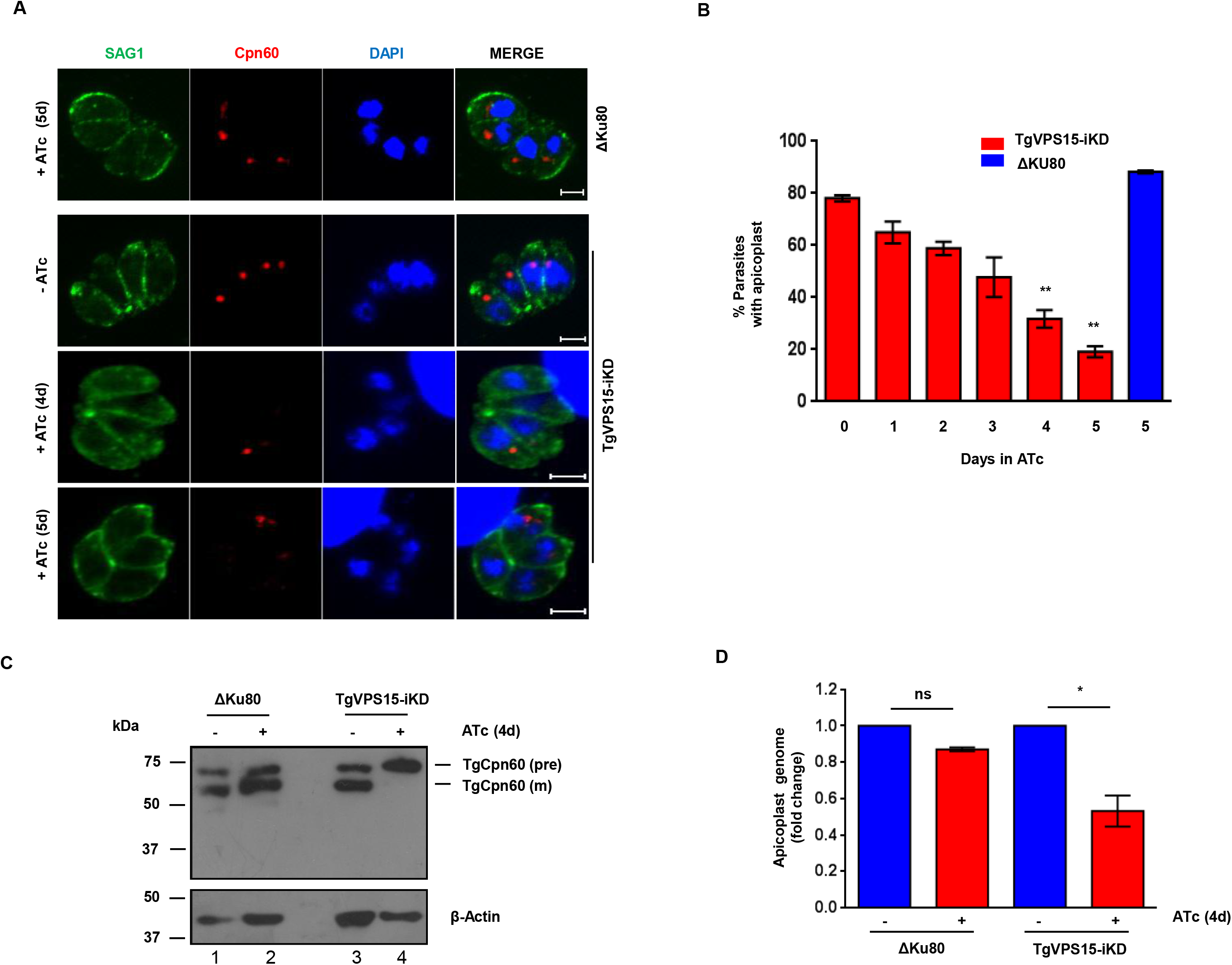
TgVPS15 regulates apicoplast biogenesis. A. IFA was performed on ΔKu80 (image not shown here) or TgVPS15-iKD parasites treated with ATc for 1-5d to detect apicoplast using an antibody against Cpn60 and the pellicle was stained using anti-SAG1. ATc treatment of TgVPS15-iKD parasites resulted in a loss of apicoplast. Representative images shown here are for experiments done after 4 and 5 days of ATc treatment. B. TgVPS15-iKD (red) or ΔKu80 (blue) parasites were treated with ATc for indicated duration as described in panel A and IFA was performed using Cpn60 antibody. % parasites with an apicoplast were counted (Mean±SE, n=3, ** P<0.01, ANOVA). C. ΔKu80 or TgVPS15-iKD parasites were treated with ATc for 96h or 4d. Western blotting was performed on parasite lysates using anti-Cpn60 antibody. Both mature (m) and precursor (pre) Cpn60 was detected in all cases except in ATc-treated TgVPS15-iKD parasites, which exhibited mainly the precursor Cpn60 form. D. Quantitative PCR was performed using total DNA from ΔKu80 or TgVPS15-iKD parasites treated with ATc for 96h or 4d and primers specific to nuclear and apicoplast coded genes. Fold change in genomic equivalents for apicoplast genome which was normalized with respect to the nuclear genome is provided [Mean± SE, n=3, * P<0.05, ANOVA].

### TgVPS15 may regulate apicoplast biogenesis via TgATG8

The ubiquitin like protein ATG8 plays a central role in autophagy; it is recruited to the autophagosome membrane upon induction of autophagy (Besteiro et al, 2011; Kong-Hap et al, 2013; Leveque et al, 2015). Interestingly, ATG8 is localized to the apicoplast outer membrane, conjugated to phosphatidyl-ethanolamine (PE) and is involved in apicoplast inheritance in *Plasmodium* and *Toxoplasma* parasites (Leveque et al, 2015). Furthermore, PI3K/PI3P is critical for TgATG8 localization to the apicoplast (Bansal et al, 2017). Therefore, it was worth testing if TgVPS15 regulates TgATG8. To this end, a parasite line was generated in which GFP-TgATG8 was integrated for ectopic expression in TgVPS15-iKD parasites. TgATG8 was found at the apicoplast and also in the cytoplasm as described previously (Bansal et al, 2017). Upon TgVPS15 depletion, TgATG8 was mainly diffused in the cytoplasm in a significant number of parasites (Fig. 4A). Since a significant number of parasites (~70-80%, Fig. 3A,B) had lost the apicoplast upon TgVPS15 depletion, the TgATG8 localization in remaining parasites-that possessed the apicoplast-was determined. More than half of these parasites did not exhibit TgATG8 at the apicoplast (Fig. 4A). In order to be functionally active, TgATG8 needs to be conjugated to phosphatidylethanolamine (PE), which is critical for its localization at the apicoplast membrane and subsequent segregation.

**Figure 4.**
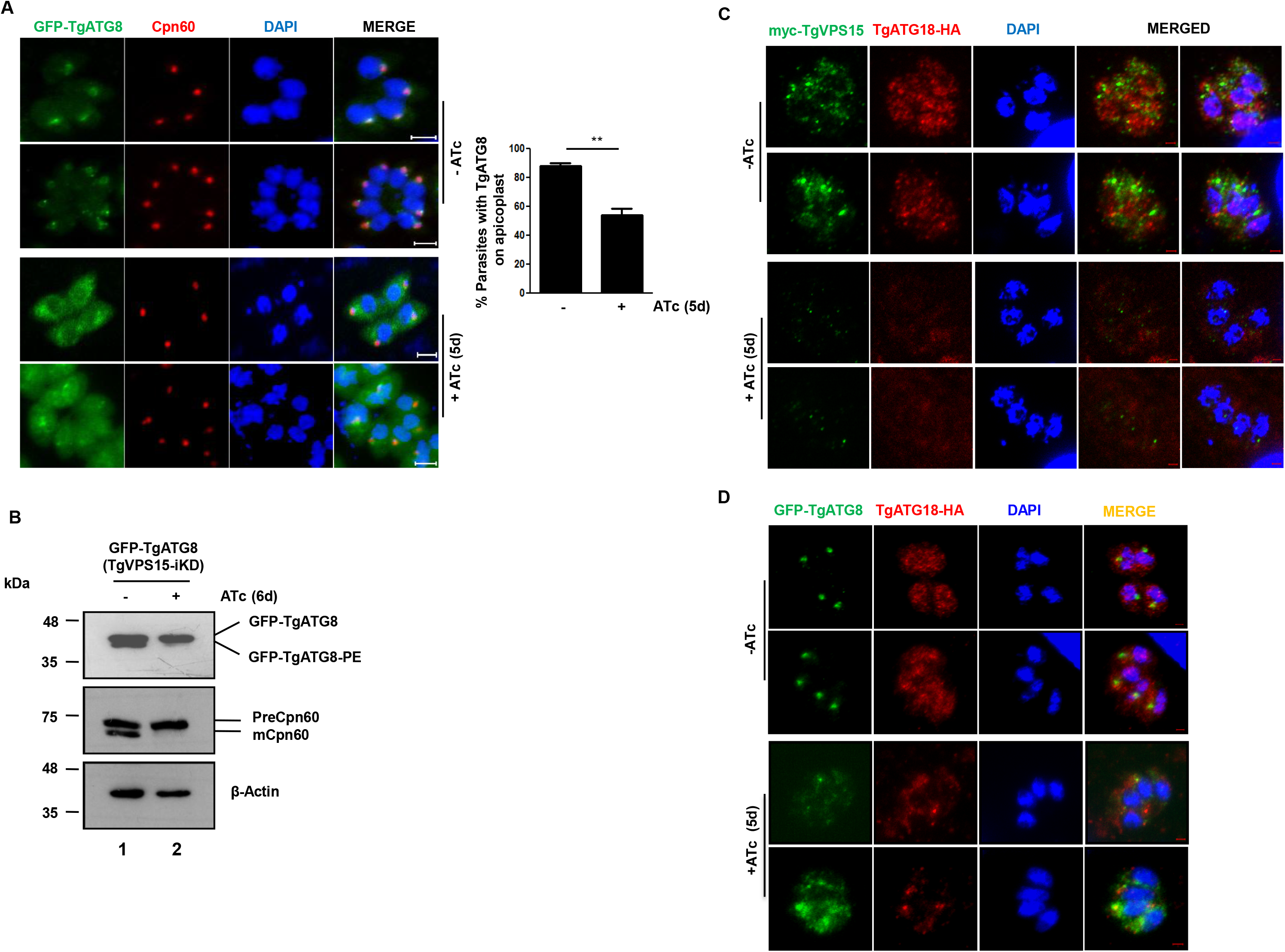
Regulation of TgATG8 and TgATG18 by TgVPS15. A. TgVPS15-iKD parasites expressing GFP-TgATG8 were treated with ATc for 5d. IFA was performed to detect apicoplast protein Cpn60 and GFP fluorescence was used to localize TgATG8. TgATG8 co-localized with the apicoplast and was also observed in the cytoplasm of untreated parasites. It was predominantly present in the cytoplasm of parasites and was largely absent from the apicoplast upon ATc treatment. *Bottom Panel,* % parasites in which TgATG8 was localized to the apicoplast (mean± SE, n=3, t-test, **, P<0.01). Please note that only those parasites that possessed a apicopalst as indicated by Cpn60 staining were used for quantitation. B. TgVPS15-iKD parasites expressing GFP-TgATG8 were treated with ATc. Subsequently, parasite lysates were electrophoresed using urea-SDS PAGE followed by Western blotting using anti-GFP antibody to detect unmodified or PE conjugated form of TgATG8 and anti-Cpn60 antibody which detected precursor or mature Cpn60. Actin was used as a loading control. C. TgVPS15-iKD/ TgATG18-HA parasites in which TgVPS15 and TgATG18 are myc and HA-tagged respectively were treated with ATc for 5d. IFA was performed using anti-myc and anti-HA antibodies followed by confocal microscopy. ATc-treatment impaired the punctate and vesicular structures of TgATG 18 (-ATc) as it was less punctate and more diffuse in the cytoplasm (+ATc). The brightness of red channel in images in +ATc condition is increased for better visualization. D. TgVPS15-iKD/ TgATG18-HA/GFP-TgATG8 parasites expressing GFP-TgATG8 and in which TgATG18 was HA-tagged were treated with ATc for 5d. IFA was performed using anti-HA antibody and anti-GFP antibody. Maximum Intensity Projection (MIP) is provided which revealed that TgATG18 was present in punctate vesicular structures but ATc treatment retained it mainly in the cytoplasm.

TgATG8 exhibited differential mobility on SDS-PAGE/Western blotting as a result of its PE conjugation, which links it to the apicoplast membrane. TgVPS15 depleted parasites exhibited mainly the unconjugated form of TgATG8 (Fig. 4B) further confirming its absence from the apicoplast. Collectively, these data suggested that TgVPS15 regulates PI3P formation. In turn, PI3P may regulate the localization of TgATG8 to the apicoplast and conjugation to its membrane (Kong-Hap et al, 2013; Leveque et al, 2015) via its downstream effectors.

TgATG18 (TGGT1_220160), which has also been referred to as TgPROP2 (Nguyen et al, 2018), was previously identified as a PI3P and PI(3,5)P2 interacting protein in *Toxoplasma,* and is involved in apicoplast biogenesis. The localization of TgATG18 to vesicular compartments is dependent on its interaction with these 3’-PIPs and is critical for its function (Bansal et al, 2017). In addition, TgATG18 was shown to regulate TgATG8 localization to the apicoplast. Given TgVPS15 facilitates the formation of PI3P in the parasite, we explored if it regulates TgATG18 by tagging it at its C-terminus in TgVPS15-iKD parasites (TgVPS15-iKD/TgATG18). As reported previously, TgATG18 localized to punctate vesicle-like structures in parasite cytoplasm. TgVPS15 also exhibited punctate vesicular localization. However, co-localization between these proteins was at best observed only occasionally (Fig. 4C).

Next, the effect of TgVPS15 depletion on TgATG18 was assessed in both TgVPS15-iKD/TgATG18 and TgVPS15-iKD/GFPATG8/TgATG18 parasites. ATc addition caused a significant reduction in TgATG18-positive puncta or vesicles and it appeared to be more diffuse in both these parasite lines (Fig. 4C-D) indicating that TgVPS15 regulates the localization of TgATG18 to vesicular organelles. Since previous studies indicated that TgATG18 localizes to vesicles via association with 3’-PIPs, (Bansal et al, 2017), present findings support the notion that TgVPS15 may regulate TgATG18 localization by its ability to promote PI3P formation via TgPI3K. TgATG18, in turn, may regulate the trafficking of TgATG8 to the apicoplast, which is important for apicoplast biogenesis (Bansal et al, 2017).

### TgVPS15 regulates autophagy in the parasite

PI3P is formed by PI3-Kinase/VPS34 activity and plays a key role in autophagy in yeast and mammals. VPS15 is known to regulate autophagy as it regulates PI3P formation via PI3K/VPS34 in yeast and mammals (Burman & Ktistakis, 2010). *Toxoplasma* is known to undergo autophagy, which is indicated by the formation of TgATG8 positive autophagosome upon nutrient deprivation (Besteiro et al, 2011; Kong-Hap et al, 2013). Therefore, it was pertinent to study the role of TgVPS15 in autophagy and TgVPS15-iKD/GFP-TgATG8 parasite line was used for this purpose. GFP-ATG8 was mainly diffuse in parasite cytoplasm in extracellular tachyzoites in complete culture medium. Upon nutrient deprivation in HBSS for 8h, there was a dramatic change in TgATG8 localization as it was localized to punctate structures, which represented autophagosomes (Fig. 5A), as reported previously (Besteiro et al, 2011; Kong-Hap et al, 2013). TgVPS15 depletion resulted in a marked reduction in parasites with GFP-TgATG8 positive puncta suggesting impaired autophagosome formation (Fig. 5A). TgATG8 is conjugated to PE on autophagosome membrane (Besteiro et al, 2011), which was enhanced under starvation condition as suggested by Western blotting (Fig. 5B, lane 2). However, TgVPS15 depletion caused a significant reduction in this form, which exhibited faster mobility on Urea-PAGE and was consistent with the loss of autophgosomes (Fig. 5B). These data suggested that TgVPS15 regulates autophagy induced by nutrient deprivation in *Toxoplasma gondii*.

**Figure 5.**
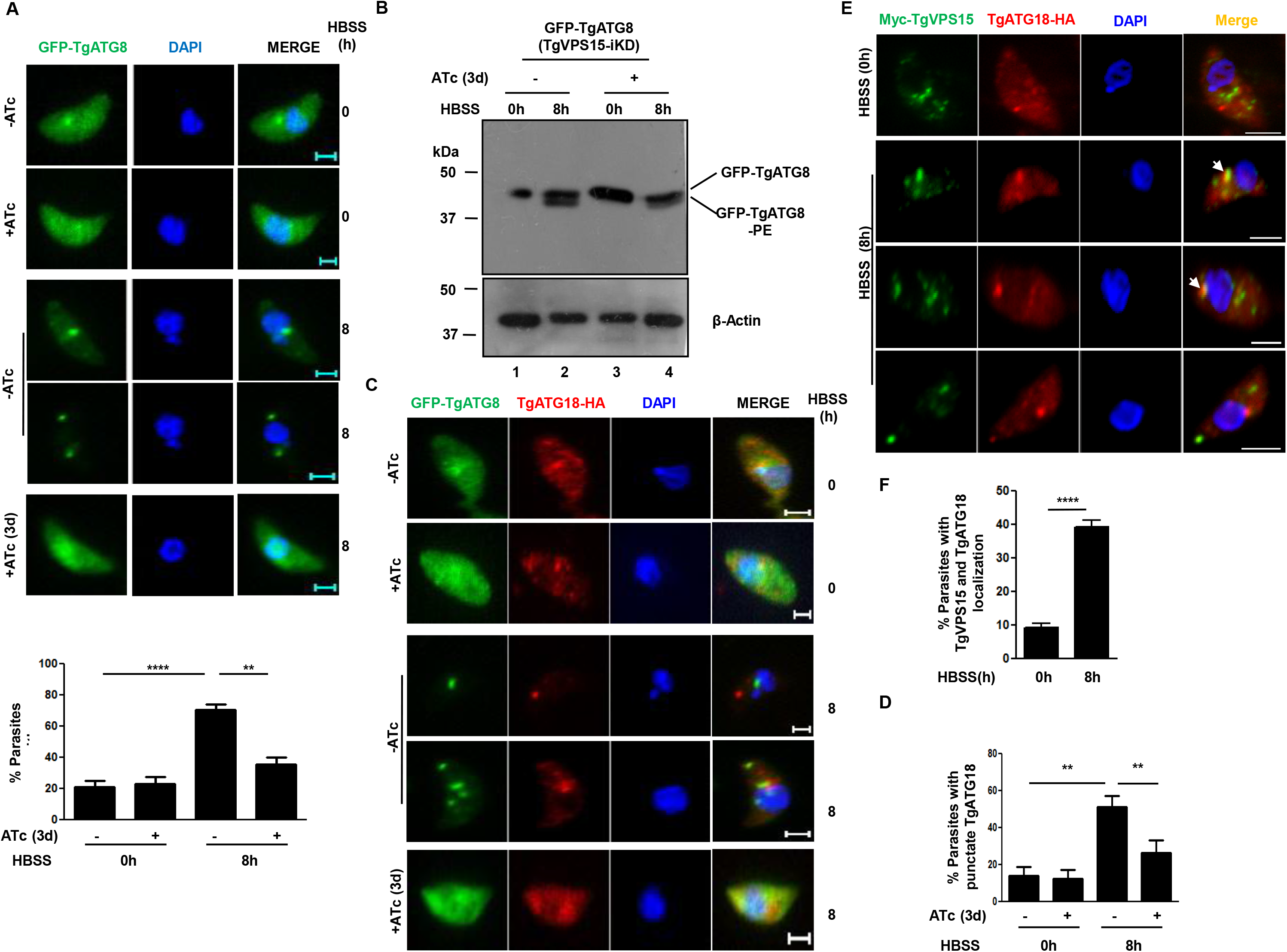
TgVPS15 regulates autophagy in *Toxoplasma*. A. TgVPS15/GFP-TgATG8 tachyzoites were left untreated or treated with ATc for 2d. Subsequently, extracellular tachyzoites were incubated in complete medium (0h) or HBSS (8h). Parasites were fixed and analyzed by fluorescence microscopy. GFP-ATG8 was mainly cytoplasmic but upon starvation in HBSS it labelled autophagosomes. Upon ATc addition to parasites in HBSS medium these punctii/autophagosomes were lost in a significant number of parasites, which were quantitated by counting at least 50 parasites (lower panel). Data represent Mean ± SE, n=3, ANOVA, ****, P<0.0001, **, P<0.01. B. Autophagy was induced in TgVPS15/GFP-TgATG8 tachyzoites as described in A. Subsequently, parasite lysates were electrophoresed by using urea-SDS PAGE followed by Western blotting using anti-GFP antibody to detect unmodified or PE conjugated form of TgATG8. Actin was used as a loading control. C-D. TgVPS15-iKD/TgATG18-HA/GFP-TgATG8 parasites were left untreated or treated with ATc for 2d. Subsequently, extracellular tachyzoites were incubated complete medium (0h) or HBSS (8h). Parasites were fixed to perform IFA using anti-HA antibody, which revealed TgATG18-HA localization to larger puncta observed in HBSS was lost upon addition of ATc. TgATG18-HA puncta were quantitated by counting at least 50 parasites (Panel D). Data represents Mean± SE, n=3, ANOVA, **, P<0.01. E. Myc-TgVPS15-iKD/TgATG18-HA extracellular tachyzoites were incubated in complete medium (0h) or HBSS (8h). Subsequently, parasites were fixed to perform IFA using anti-HA and anti-myc antibody, which revealed that TgATG18-HA as well as TgVPS15 localization changed to a larger puncta in HBSS. In some cases, they co-localized to these larger puncta (arrows). F. Quantitation of parasites in which TgATG18 and TgVPS15 co-localized by counting at least 50 parasites (panel E). Data represents Mean± SE, n=3, ANOVA, ****, P<0.0001.

### TgVPS15 mediated regulation of TgATG18 is important for autophagy

As mentioned above, the interaction of TgATG18 with PI3P and PI(3,5)P2 has been demonstrated (Bansal et al, 2017). However, its role in autophagy has remained unknown. Since TgVPS15 regulates PI3P formation, we tested the involvement of TgATG18 in autophagy. For this purpose, TgATG18 was HA-tagged at the C-terminus in TgVPS15-iKD line that expressed GFP-TgATG8 ectopically. IFA studies revealed that the localization of TgATG18 changed from small punctii to large punctate structures under starvation (Fig. 5C, 5E) but co-localization with TgATG8 on these puncta was observed in only a very few parasites (Fig. 5C, 6B, ~10% parasites). At best, TgATG18 and TgATG8 appeared to be more proximal under nutrient limiting conditions (Fig. 5C, 6B). It is likely that TgATG18 localizes to early or pre-autophagosome compartment, which was also suggested in a previous study in which TgATG18 co-localized with TgATG9 in puncta in starvation conditions (Nguyen et al, 2018). TgVPS15 depletion resulted in a dramatic loss of TgATG18-puncta under starvation (Fig. 5C, 5D), which correlated well with the loss of the autophagosomes. Next, we tested if TgVPS15 co-localized under starvation conditions and used TgVPS15-iKD/TgATG18-HA parasite line for this purpose. While TgVPS15 and TgATG18 were in distinct locations in complete medium, nutrient deprivation resulted in formation of bigger puncta with TgVPS15 staining, which was also the case with TgATG18. Interestingly, in almost 40% of parasites TgVPS15 and TgATG18 co-localized in these puncta, which as described above may be a pre-autophagosome compartment (Fig. 5E,F).

**Figure 6.**
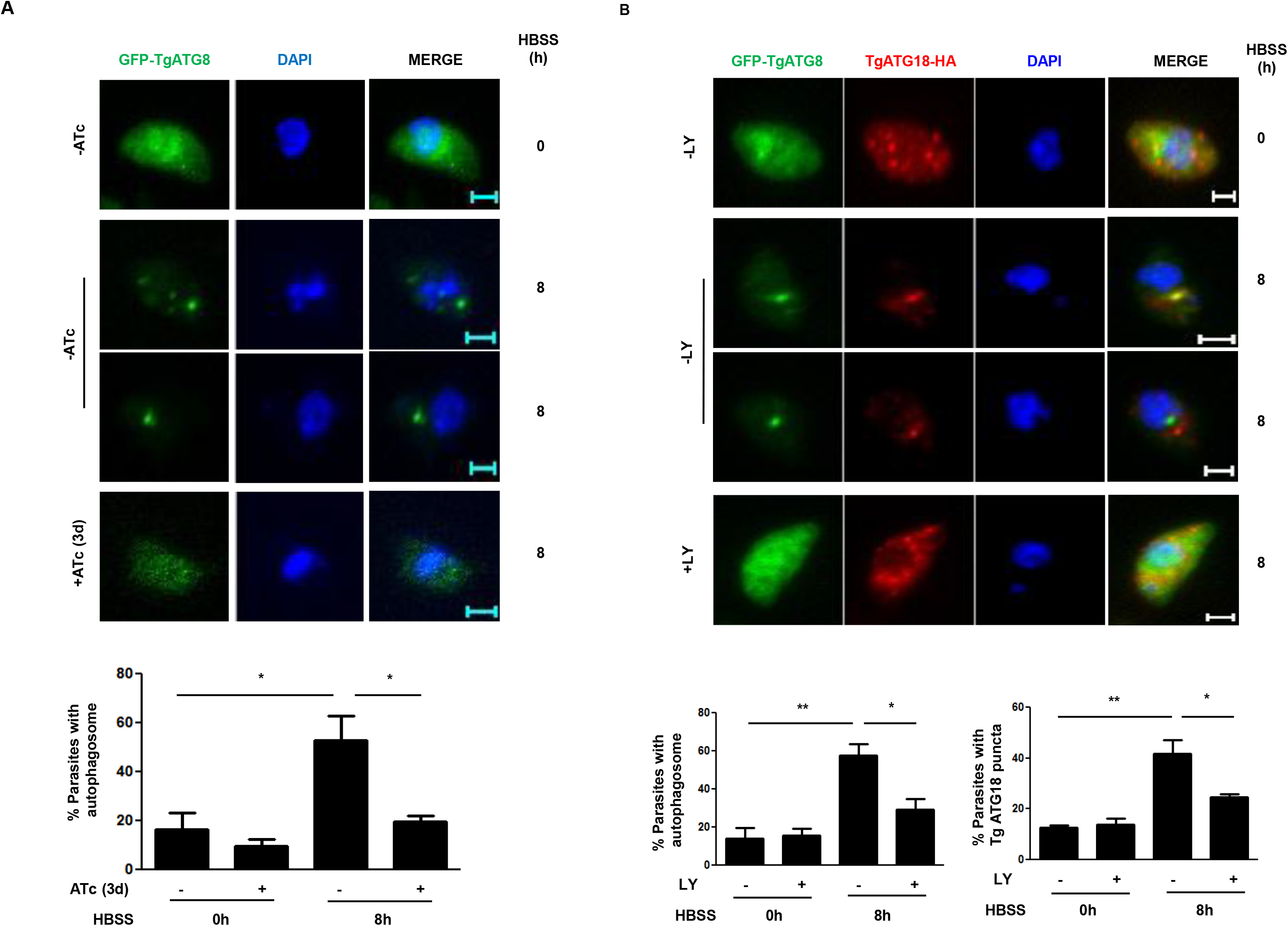
TgPI3K and TgATG18 regulate autophagy. A. TgATG18-iKD/GFP-ATG8 parasites were treated with ATc and extracellular tachyzoites were either cultured in complete medium (0h) or HBSS (8h) medium. Subsequently, parasites were fixed and analyzed by fluorescence microscopy. Upon ATc addition to parasites in HBSS medium, GFP-ATG8 positive autophagosomes were lost in a significant number of parasites, which were quantitated by counting at least 50 parasites (bottom panel). Data represents Mean± SE (n=3, ANOVA, *, P<0.05). B. TgATG18-iKD/GFP-ATG8 extracellular tachyzoites were treated with DMSO (-) or 100 μM LY294002 and cultured in complete medium (0h) or HBSS (8h). Subsequently, parasites were fixed and analyzed by fluorescence microscopy. The number of parasites with TgATG18 or TgATG8 positive puncta was counted under each condition (bottom panel). Data represents Mean± SE (n=3, ANOVA, **, P<0.01, *, P<0.05).

In order to probe if there is a direct-link between TgATG18 and autophagy, we tested if TgATG18 regulates TgATG8 during autophagy. To this end, a previously reported TgATG18-iKD/GFP-ATG8 parasite line was used (Bansal et al, 2017). TgATG8 positive autophagosomes formed under starvation were significantly impaired upon TgATG18 depletion (Fig. 6A) suggesting that TgATG18 promotes autophagosome formation in the parasite.

Even though most of the above-mentioned functions of TgVPS15 may be attributed to its ability to regulate PI3P formation in the parasite, direct involvement of PI3K remained to be tested which was done by using a PI3K-inhibitor LY294002 that can inhibit PI3P production in *Toxoplasma* by inhibiting TgPI3K (Tawk et al, 2011). LY294002 blocked the TgATG8-positive autophagosomes that are formed under starvation conditions. In addition, TgATG18-positive puncta were also reduced significantly by this inhibitor (Fig. 6B), which could be explained by the fact that TgATG18-interacts with PI3P (Bansal et al, 2017). Collectively, these data established that TgVPS15 regulates autophagy via its ability to regulate the formation of PI3P, which in turn regulates TgATG18 and these events are critical for TgATG8 localization to the autophagosome and the formation of this organelle (Fig. 8).

### Catalytic activity of TgVPS15 may play a role in its function

As mentioned above (Fig. 1), several key elements present in proteins kinases are either absent from TgVPS15 or are divergent. Therefore, it may be a pseudokinase like ScVPS15. However, it is known that ScVPS15 autophosphorylates and some of the key residues may be important for its function (Herman et al, 1991). We investigated if TgVPS15 activity is important for its parasitic functions. Our attempts to perform kinase assays with the kinase domain of TgVPS15 (TgVPS15-KD) in *E.Coli* were inconsistent as expression of the recombinant was poor. Therefore, we used an independent approach to address this issue by ectopically expressing either WT TgVPS15 or two of its mutants. These mutants were generated by replacing D216 with A, which is part of the divergent HGD motif and may act as a catalytic base (Nolen et al, 2004). A E268A mutant of the conserved APE motif, which is important for key interactions necessary for stabilization of the activation loop (Nolen et al, 2004), was also made. D216A and E268A mutants were ectopically expressed in TgVPS15-iKD with a Ty-tag as described above for the WT TgVPS15 complementation line (TgVPS15-iC^WT^) (Supp. Fig. S5).

First, the role of these residues in the ability of TgVPS15 to promote PI3P was assessed. For this purpose, the 2xFYVE reporter construct described above (Fig. 2G) was expressed in these parasite lines. The loss of PI3P at the apicoplast in TgVPS15-iKD parasites upon ATc addition (~15% parasites with PI3P at apicoplast) was efficiently reverted in TgVPS15-iC parasites (~44%) in which WT TgVPS15 were complemented. In contrast, both D216A and E264A mutants (~15%) were unable to restore PI3P formation efficiently. Based on these data, it is reasonable to conclude that TgVPS15 catalytic activity may be critical for the regulation of TgPI3K and promote the formation of PI3P in the parasite.

Next, the role of TgVPS15 activity on its parasitic functions was investigated. As described above, the complementation with WT TgVPS15 prevented any defects in parasite replication that were observed in TgVPS15-iKD parasites upon ATc addition. However, the expression of D216A as well as E268A mutants of TgVPS15 did not prevent the arrest in parasite replication (Fig. 7B). Next, we assessed apicoplast biogenesis in these parasite lines. As reported above (Fig. 3A-B), only ~20% of ATc treated TgVPS15-iKD parasites exhibited the presence of apicoplast, which significantly increased (~40%) when WT TgVPS15 was ectopically expressed (TgVPS15-iC^WT^). In contrast, parasites expressing the D216A and E268A in TgVPS15-iKD parasites also exhibited only ~20% parasites with an apicoplast (Fig. 7C, Supp. Fig. S6). These data suggested that D216 and E268 were critical for TgVPS15 function in parasite replication, which is dependent on its ability to regulate apicoplast biogenesis.

**Figure 7.**
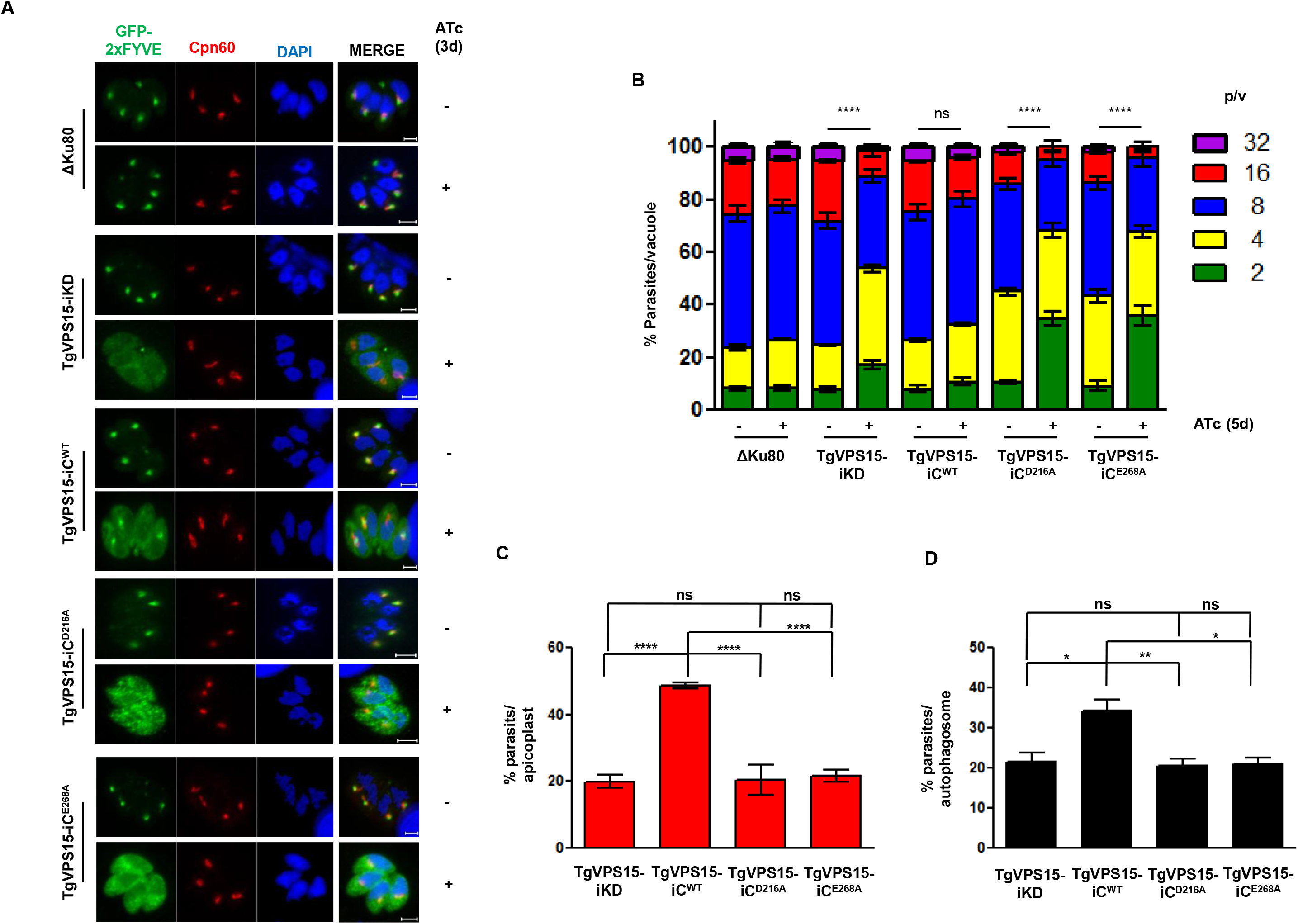
Role of catalytic site residues in the TgVPS15 function. A. DD-GFP-2xFYVE was expressed in TgVPS15-iKD parasites or these parasites complemented with a copy of Ty-tagged WT TgVPS15, or its D216A or E268A mutants. Subsequently, parasites were treated with ATc for 72h (3d) or left untreated. Prior to fixation, Shield-1 was added for 30 minutes to stabilize the expression of GFP-2xFYVE. IFA was performed using anti-Cpn60 antibody. The image of only those parasites in which apicoplast was present is shown. While WT TgVPS15 complementation restored PI3P formation indicated by 2xFYVE staining at the apicoplast, majority of parasites complemented with D216A and E268A exhibited diffuse 2xFYVE in the cytoplasm. B. TgVPS15-iKD parasites were complemented with a copy of Ty-tagged WT TgVPS15 or its D216A and E268A mutants. The indicated parasite lines were preincubated for 96h or 4d in culture medium in the presence (+) or absence (-) of ATc and were subsequently allowed to invade fresh HFFs in the presence or absence of ATc. The number of parasites per vacuole was determined after 24h. Data represent mean ± SE, n=3 and at least 200 vacuoles were counted for each condition (n=3, ****, P<0.0001, ANOVA, 8p/v, ns-not significant). C. The indicated parasite lines were pre-treated with ATc for 4d and IFA was performed using Cpn60 antibody after additional 24h (Supp. Fig. S6). % parasites containing the apicoplast was determined by counting parasites in IFA images (Supp. Fig. S6). Mean±SE, n=3, ** P<0.01, ANOVA, ns-not significant. D. GFP-ATG8 was ectopically expressed in indicated parasite lines. Parasites were treated with ATc for 2d and extracellular tachyzoites were either cultured in complete (0h) or HBSS medium (8h). Subsequently, parasites were fixed and analyzed by fluorescence microscopy (Supp. Fig. S7). GFP-ATG8 positive autophagosomes were counted from at least 50 parasites and % parasites possessing autophagosomes was determined (Mean±SE, n=3,* P<0.05, **P<0.01, ANOVA, ns-not significant).

We also investigated the involvement of these TgVPS15 active site residues in autophagy by using the above mentioned parasite lines. A significant increase in autophagosome formation was observed in TgVPS15-iC parasites was observed when compared to ATc-treated TgVPS15-iKD parasites. In contrast, the expression of both D216A and E268A mutants did not cause a similar increase. These data suggested that catalytic site residues play a critical role in autophagy as well (Fig. 7D, Supp. Fig. S7) and hint at the possible involvement of TgVPS15 activity even though it is not a conventional kinase.

## Discussion

3’-PIPs are involved in critical cellular processes like membrane and protein trafficking, organelle homeostasis, autophagy and signalling (Balla, 2013). These PIPs are critical for the development of apicomplexan parasites *Plasmodium falciparum* and *Toxoplasma gondii* (Cernikova et al, 2019). PI3P is important for apicoplast biogenesis in both *Plasmodium* and *Toxoplasma* (Bansal et al, 2017; Tawk et al, 2010; Tawk et al, 2011). In addition, it is involved in the trafficking of haemoglobin to the digestive vacuole of *Plasmodium* (McIntosh et al, 2007; Vaid et al, 2010) and is implicated in resistance to artemisinin (Mbengue et al, 2015). A single VPS34-like PI3-kinase is involved in the biogenesis of 3’-PIPs is indispensable for these parasites and depletion of PI3P impairs parasite growth (Daher et al, 2015; Tawk et al, 2010; Tawk et al, 2011; Vaid et al, 2010). However, the mechanism via which PI3Ks and PI3P regulate the development of the parasite have remained largely unknown. In addition, a putative PIKfyve kinase homologue, which is likely to be involved in PI(3,5)P2 formation is coded only in *Toxoplasma* and is also involved in apicoplast homoeostasis (Daher et al, 2015). In the present study, we demonstrate that TgVPS15 is a regulator of TgPI3K activity as its abrogation results in depletion of PI3P from the parasite (Fig. 2G). Although biochemical characterization of TgPIKfyve has not been done, it is likely that it may use PI3P as a substrate to convert it to PI(3,5)P2 like the yeast enzyme. Therefore, it is very likely that TgVPS15 may also regulate PI(3,5)P2 generation as PI3P serves as a precursor for PI(3,5)P2 formation, which we could not test in the present study. Typically, VPS15 from most organisms is classified as a pseudokinase due to the following reasons: a. several key motifs that are absent from TgVPS15 kinase domain when compared to typical protein kinases (Fig. 1B). For example, the glycine rich loop, HRD and DFG motifs are either absent or are diverse (Herman et al, 1991; Rostislavleva et al, 2015). TgVPS15 also exhibits similar differences (Fig. 1).

b. There is no strong evidence for VPS15 acting as a protein kinase as its direct substrates have not been reported and only autophosphorylation has been reported for ScVPS15 (Herman et al, 1991; Stack & Emr, 1994). Interestingly, mutation of critical kinase domain residues postulated to play a role in its catalytic activity prevent its autophosphorylation, PI3P generation, impair its function in endosome sorting and autophagy (Herman et al, 1991; Stack et al, 1995; Stack et al, 1993).

Due to the inability to express and purify the recombinant TgVPS15 kinase domain efficiently, we could not perform kinase activity assays to ascertain if it possesses kinase activity. However, the mutation of D216 and E268 in critical region-HGD and APE motifs-of the kinase domain prevented the rescue of parasite growth, apicoplast inheritance and autophagy when TgVPS15 was depleted, which could be attributed to impaired PI3P formation (Fig. 7A). These data strongly suggested that despite possessing features of pseudokinase, TgVPS15 activity contributes to its function. These results exhibited similarity with the observations made in the case of ScVPS15, in which similar mutations also impaired its cellular functions in the yeast (Herman et al, 1991). It is possible that TgVPS15 may be autophosphorylated, which may promote its association with TgVPS34 and other regulatory partners, which has been reported for ScVPS15 (Stack et al, 1995).

TgVPS15 is present in vesicular structures in the parasite which may be suggestive of its role in membrane/protein trafficking (Fig. 2B, 4C). Its depletion resulted in a loss of the apicoplast (Fig. 3) and explained the defects in parasite replication observed after the first replication cycle (Fig. 2E), which is typical of “delayed death” observed due to the loss of the apicoplast (Roos et al, 1999). Our studies identify TgVPS15 to be an upstream regulator of TgPI3K via which PI3P is generated, which results in the regulation of TgATG18, which has been previously demonstrated to play a role in apicoplast biogenesis (Bansal et al, 2017). The interaction of TgATG18 with 3’-PIPs is critical for its cellular localization and function (Bansal et al, 2017), which is regulated by TgVPS15 (Fig 4D). Given TgATG18 regulates the localization of TgATG8 to the apicoplast membrane (Bansal et al, 2017), the TgVPS15 directed TgATG8 apicoplast localization (Fig. 4A) is most likely due to its ability to regulate TgATG18 (Fig. 4C,D). It remains unclear how TgATG18 regulates the trafficking of TgATG8 to the apicoplast. Since they do not co-localize, other molecules like the components of trafficking machinery may be involved in this process, which need to be identified (Bansal et al, 2017). Previous studies have suggested that TgATG8 may facilitate an apicoplast-centrosome link (Leveque et al, 2015), which allows it to play a role in parasite division.

VPS15 associates with PI3-kinase VPS34 in two different complexes and promotes its activation. Complex I comprises of ATG14, which further strengthens the association between VPS34-VPS15 along with ATG6 and in complex II VPS38 replaces ATG14 (Kim et al, 2013; Ohashi et al, 2019; Rostislavleva et al, 2015). Typically, complex I predominantly regulates autophagy whereas Complex II is involved in vacuolar protein sorting ((Itakura et al, 2008)). Out of the VPS34 complex proteins only VPS15, VPS34 and ATG6 homologues have been identified in *Toxoplasma* and ATG6 seems to be absent in *Plasmodium* (Sakamoto et al, 2021). These observations hint at a distinct mechanism of regulation of VPS34 complex in these parasites. Despite these differences, present studies indicated that TgVPS15 regulates the formation of PI3P via TgPI3K regulation in the parasite. The localization of TgATG18 changes from small vesicles to a bigger puncta when parasites are starved for nutrients (Fig. 5C). Previously, it was co-localized with pre-autophagosomal protein TgATG9 (Nguyen et al, 2018), which suggests that TgATG18 may translocate to pre-autophagosomal compartment upon nutrient deprivation. TgVPS15, which also resides in vesicles under steady state conditions, exhibited a change in its localization to bigger puncta upon nutrient deprivation (Fig. 5E) and co-localized with TgATG18 in these puncta likely to be pre-autophagosomal compartments. TgVPS15 depletion impaired the targeting of TgATG18 to this pre-autophagosome like organelle (Fig. 5C), which was supported by the fact that it interacts with PI3P (Bansal et al, 2017). Subsequently, TgATG18 regulates TgATG8 localization to the autophagosome (Fig. 6A) and formation of this organelle, TgVPS15 depletion (Fig. 5A and C) as well as pharmacological inhibition of PI3K (Fig. 6B) both which impair PI3P formation caused similar defects in TgATG18 and TgATG8 localization..

Present findings highlight the importance of PI3P generation by TgVPS15 which it may achieve by regulating TgPI3K either by facilitating its interaction with other important proteins and/or regulating its activity directly. PI3P formed regulates trafficking of TgATG18 to preautophagsome, which in-turn regulates the localization and conjugation of TgATG8 to autophagosome membrane where is conjugated to PE and promotes autophagosome formation and autophagy (Fig. 8).

**Figure 8.**
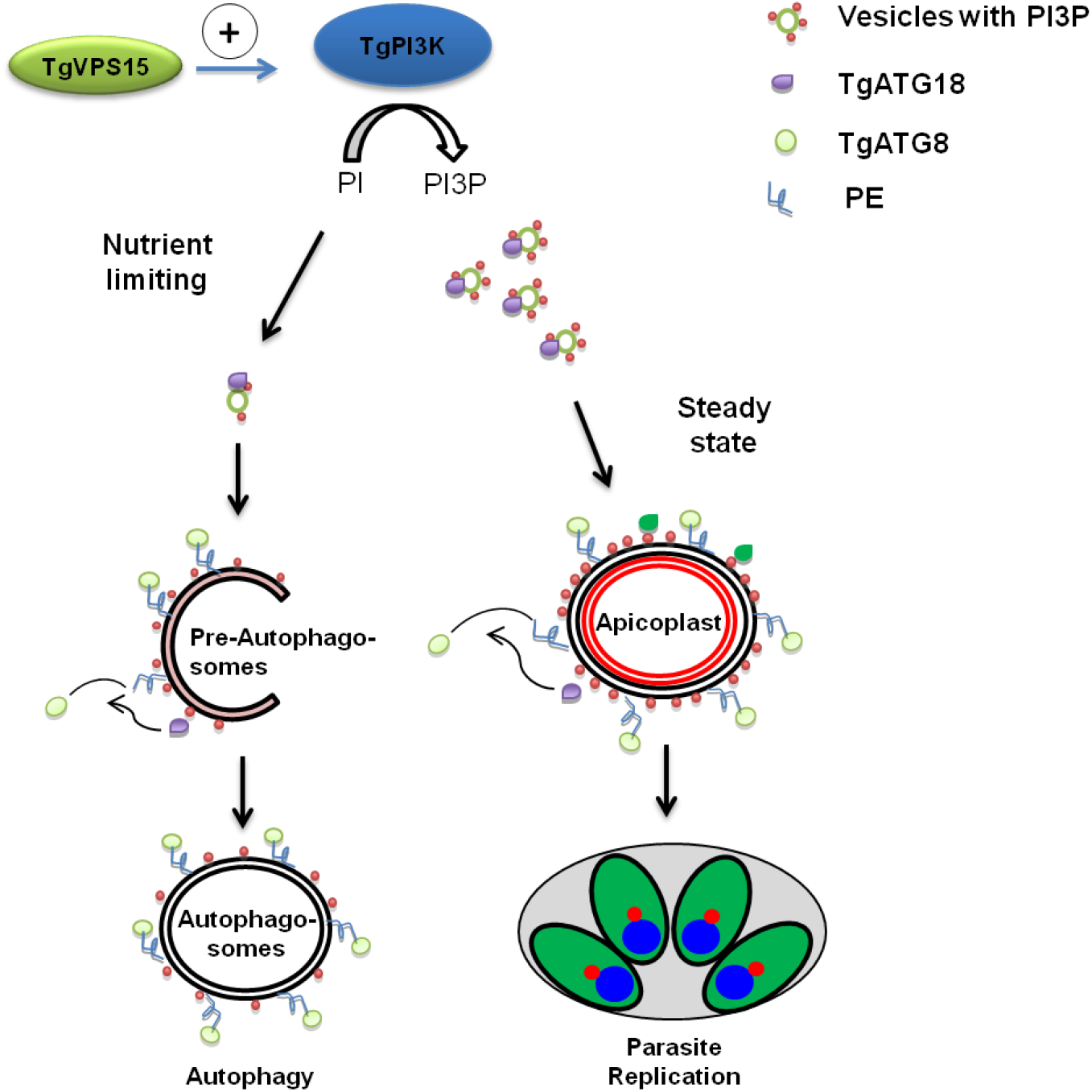
A signaling pathway involving TgVPS15 regulates apicoplast inheritance and autophagy. TgVPS15 may regulate TgPI3K and facilitate the formation of PI3P (Fig. 2B). Under steady state conditions, PI3P regulates the localization of TgATG18 to vesicular compartments (Fig. 4C and D), which in turn promotes the trafficking of TgATG8 to the apicoplast membrane (Bansal et al, 2017) and is critical for the inheritance of the apicoplast during parasite division. Under nutrient limiting conditions, TgVPS15 may facilitate the generation of PI3P via TgPI3K at preautophagosome-like vesicles (Fig. 5E) resulting in the localization of TgATG18 to this compartment (Fig. 5C and E), which sets the stage for the formation of autophagosome to which TgATG8 is trafficked and where it is conjugated to autophagosomal membrane (Fig. 5A and 6A) and contributes to the process of autophagy.

These studies unravel a dual role of TgVPS15 in *Toxoplasma gondii* and both are dependent on its ability to facilitate PI3P formation; it regulates apicoplast inheritance during parasite lytic cycle and autophagy under nutrient-limiting conditions (Fig. 8).

## Materials and Methods

### *Toxoplasma* cultures, plaque and growth rate assays

*Toxoplasma gondii* tachyzoites RHΔhxgprt (Donald et al, 1996) and RHΔhxgprtΔKu80 (Huynh & Carruthers, 2009) were grown in human foreskin fibroblast HFFs and maintained in Dulbecco’s modified Eagle’s medium supplemented with 10% fetal bovine serum and 2mM glutamine at 37°C, 5% CO_2_ in humidified incubator. 500nM of Shield-1 (Shld-1) was added for destabilization domain (DD) fusion proteins, 1μg/ml of anhydrotetracycline (ATc) was used for TgVPS15-iKD and TgATG18-iKD parasites.

#### *T. gondii* plaque and intracellular growth assays

For plaque assays, parasites were pretreated for 48h or 2d with 1μg/ml ATc. Freshly egressed parasites were normalized and used to re-infect HFF monolayer and treated with ATc for ~ 7 days. The cells were fixed with icecold methanol and stained with crystal violet. The intracellular growth assay of TgVPS15-iKD parasites was performed by pre-treatment with ATc for 48 or 96h unless indicated otherwise, and the parasites were cultured further for specified time in the presence or absence of ATc. IFA was performed after PFA fixation using anti-GAP45, anti-SAG1 and anti-Cpn60 antibody. The number of parasites per vacuole was determined and ~ 200 vacuoles were counted for each condition.

### Generation of transgenic parasite lines

PCR primers used for generating various constructs are listed in Table S1.

#### TgVPS15-iKD

A 2519 bp fragment corresponding to the 3’ homology arm from *TgVPS15* was amplified using primers 3/4 (for N-myc tag) and 3”/4 (without tag) by PCR from genomic DNA and subcloned using BglII and NotI (for N-myc tagging) and EcoRV and NotI (without N-myc tagging) sites of TATi-HXGPRT-tetO7Sag1MycTag vector (Salamun et al, 2014). The 5’-homology arm flanking untranslated region (UTR) upstream of the *TgVPS15* promoter (2,625bp) was PCR amplified by using primers 1/2 and cloned using NcoI and BamHI sites in TATi-HXGPRT-tetO7Sag1-MycNtTgVPS15 plasmid construct. The resulting plasmid pVPS15-TATi-HXGPRT-tetO7Sag1-*TgVPS15* was transfected in ΔKu80 parasites by electroporation of 50μg of this construct (linearized by NcoI and NotI digestion prior to transfection). Parasites were subsequently subjected to 25μg/ml mycophenolic acid (MPA) and 50μg/ml xanthine selection. The drug-resistant parasites were cloned by limiting dilution and possible clones were screened by PCR to detect the presence of endogenous or recombined locus.

#### TgVPS15-iKD/TgATG18-3HA

A ~1.4-kb fragment was amplified from the 3’ end of TgATG18 using *Toxoplasma gondii* genomic DNA and cloned into LIC-DHFR-3HA vector and TgVPS15-iKD parasites were electroporated with 50μg of plasmid digested with *Rsr II.* Parasites were selected on 5μM pyrimethamine and single clonal population was selected by limiting dilution cloning.

#### TgVPS15-iKD WT or mutant complemented parasites (TgVPS15-iC^WT^ TgVPS15-iC^D216A^, TgVPS15-iC^E268A^)

For complementation of TgVPS15, *TgVPS15* cDNA was synthesized (GeneArt, Invitrogen). pCTDD-HA vector (Bansal et al, 2017) (a gift from Dr. Ravikant Ranjan) was digested using *BglII* and *EcoRV* and TgVPS15 cDNA was ligated in it using Gibson assembly kit (NEB) to obtain pCT-TgVPS15-iC^WT^. Subsequently, an N-terminal Ty-tag was introduced using primers 31/32 to obtain pCT-TyTgVPS15-iC^WT^. D216A and E268A mutations were generated by site directed mutagenesis using primer 23/24 and 27/28, respectively in pET-TgVPS15_KD construct which had the kinase domain cloned in pET28A vector. Subsequently, an amplicon (I) from the kinase domain portion was amplified using primers 33/30-with an *NcoI* site in the reverse primer and using the pET construct described above containing mutant TgVPS15. Another amplicon (II) was generated using primers 29/34 from amplified from pCT-TyTgVPS15-iC^WT^ in which *NcoI* site was introduced at the 5’ end. The two fragments (I and II) were assembled using Gibson assembly kit after *NcoI* digestion of the pCT-TgVPS15-iC^WT^ described above. The replacement of the WT TgVPS15 with the desired mutant fragment from this construct was confirmed by DNA sequencing. 50μg of the circular plasmid was digested using *PmeI* (linearized) and transfected in TgVPS15-iKD and selection was done using 20μM chloramphenicol.

#### TgVPS15-iKD/GFP-TgATG8 and TgVPS15-iC^(WT/D216A/E268A^/GFP-TgATG8

GFP-TgATG8 cDNA was cloned into the pTub8-Bleo vector, which had a myc tag (a gift from Dr. Sebastien Besteiro). For stable GFP-TgATG8 expression, 50μg of vector was digested using *SacI* and a linearized copy was transfected in ΔKu80, TgVPS15-iKD or TgVPS15-iC^(WT/D216A/E268A)^ parasites. Extracellular parasites were selected with 25μg/ml phleomycin along with the other desired drugs and a clone obtained by limiting dilution was used for further studies.

#### TgVPS15-iKD/DD-GFP-2xFYVE and TgVPS15-iC^(WT/D216A/E268A)^/DD-GFP-2xFYVE

Two tandem repeats of Hrs FYVE domain (2xFYVE) were ectopically expressed as N-terminal GFP fusion protein along with a FKBP-DD domain (DDGFP-2xFYVE). For this purpose, 2xFYVE was amplified from p3E-2xFYVE_hrs, which was a gift from Rob Parton (Addgene plasmid # 67676; http://n2t.net/addgene:67676; RRID: Addgene_67676) using PCR primers 17/18 and fused to DD-GFP (amplified using primers 15/16 from pCTDD-GFP vector) using overlapping PCR followed by cloning in pCTDD-HA vector digested with *Bgl II* and *EcoRV* sites to yield pCTDD-GFP-2xFYVE-CRT construct. 50μg of *PmeI* digested pCT-DDGFP-2xFYVE-CRT plasmid was transfected in both ΔKu80 and TgVPS15-iKD parasites. Selection with chloramphenicol was done immediately after transfection and continued until the parasites appeared followed by MPA and Xanthine selection. Limiting dilution cloning was done to obtain a clone to be used for further studies.

To generate DD-GFP-2xFYVE expressing transgenic parasite in the TgVPS15-iC^(WT/D216A/E268A)^ background which as described above already contains CRT cassette. The CRT cassette in pCTDDGFP-2xFYVE-CRT was replaced with DHFR cassette. The resulting CTDDGFP-2xFYVE-DHFR subsequently obtained was transfected to generate the above mentioned lines and were kept under 5μM pyrimthamine.

#### TgVPS15-iKD_TgATG18/TgATG8

TgVPS15-iKD/TgATG18-HA parasite line described above was transfected with 50μg *SacI* digested pTub8-mycGFP-TgATG8-Bleo construct. Transfected parasites were treated with pyrimethamine and phleomycin as described and followed by limiting dilution cloning.

### Immunofluorescence assays

For immunofluorescence assays, human foreskin fibroblasts (HFFs) seeded on coverslips were infected with freshly egressed parasites. Intracellular parasites cultured for the indicated times were fixed with 4% PFA in phosphate-buffered saline (PBS) followed by incubation with various primary antibodies and processed as described previously (Bansal et al, 2021). Briefly, parasites were incubated with the relevant primary antibodies at 4°C, washed with PBS, and incubated with AlexaFluor 488/594-labeled secondary antibodies (Invitrogen) at room temperature. Following washing, either Hoechst 33342 or mounting medium containing 4-,6-diamidino-2-phenylindole (DAPI) was used to stain the nucleus. The stained parasites were examined using Axio Imager Z1 microscope or confocal microscope LSM 700 (Fig. 2C) or LSM980 (Fig. 4C) (Carl Zeiss). Z stacking during image acquisition and processing of images was done using AxioVision 4.8.2 software or ZEN 2 (blue edition). Z-stacks that best represented the immunolocalization were used for illustrations in figures unless indicated otherwise. Maximum Intensity Projection (MIP) was performed for TgATG8/TgATG18 localization as indicated in Fig. 4D.

### Immunoblotting

Freshly egressed *T. gondii* tachyzoites were used for the preparation of lysate, which was done in a buffer containing 10 mM Tris (pH 7.5), 100 mM NaCl, 5 mM EDTA, 1% Triton X100, and Complete protease inhibitor mixture (Roche Applied Science) or in 2% SDS. SDS-PAGE was performed, followed by transfer of proteins to nitrocellulose membranes. Immunoblotting was performed using various primary antibodies and antisera, and HRP labeled anti-rabbit IgG. West Pico, West Dura, or Femto enhanced chemiluminescence (ECL) substrate (Pierce) was used to develop blots following manufacturer’s instructions. For resolving PE-conjugated and unlipidated TgATG8, urea SDS-PAGE was performed prior to Western blotting (Bansal et al, 2017).

### Induction of autophagy

Extracellular parasites naturally or mechanically egressed from HFFs were washed twice in Hank’s Balanced Salt Solution (HBSS) before incubation in HBSS at 37°C for up to 8h (Kong-Hap et al, 2013). Subsequently, live imaging of GFP-TgATG8 and/or IFA was performed for detection of other proteins followed by fluorescence microscopy as described above.

### Statistical analyses

The statistical analysis was performed using Graph Pad PRISM. P-value of less than 0.05 was considered as significant and n represents the number of independent biological replicates.

## Acknowledgments

Studies were supported by grants BT/COE/34/SP15138/2015 from Department of Biotechnology and funds from NII core. P.S. is a recipient of J.C. Bose Fellowship; P.B. received Senior Research fellowship from CSIR. We appreciate reagents kindly gifted by Dr. Dominique Soldati, Dr. Sebastien Besteiro and other researchers. pCTDD-HA vector was a kind gift from Dr. Ravikant Ranjan.

## Author Contributions

PS, PB and RSR analyzed the data and wrote the manuscript. RSR performed most experiments and was supported by P.B. Figures were prepared by RSR and PS.

